# UV-Induced Keratin 1 Proteolysis Mediates UV-Induced Skin Damage

**DOI:** 10.1101/226308

**Authors:** Mingchao Zhang, Dhruba Tara Maharjan, Hao He, Yujia Li, Wei Yan, Weixin Yan, Yujie Zhu, Weihai Ying

## Abstract

Keratins play critical roles in intermediate filament formation, inflammatory responses and cellular signaling in epithelium. While keratins is a major epidermal fluorophore, the mechanisms underlying the autofluorescence (AF) of keratins and its biomedical implications have remained unknown. Our study used mouse skin as a model to study these topics, showing that UV dose-dependently induced increases in green AF at the spinous layer of the epidermis of mouse within 6 hr of the UV exposures, which may be used for non-invasive prediction of UV-induced skin damage. The UV-induced AF appears to be induced by cysteine protease-mediated keratin 1 proteolysis: 1) UV rapidly induced significant keratin 1 degradation; 2) administration of keratin 1 siRNA largely decreased the UV-induced AF; and 3) administration of E-64, a cysteine protease inhibitor, significantly attenuated the UV-induced AF and keratin 1 degradation. Our study has also suggested that the UV-induced keratin 1 proteolysis may be a novel crucial pathological factor in UV-induced skin damage, which is supported by both the findings that indicate critical biological roles of keratin 1 in epithelium and our observation that prevention of UV-induced keratin 1 proteolysis can lead to decreased UV-induced skin damage. Collectively, our study has suggested that UV-induced keratin 1 proteolysis may be a novel and valuable target for diagnosis, prevention and treatment of UV-induced skin damage.

## Introduction

Keratins play multiple significant roles in epithelium, including intermediate filament formation^1^, inflammatory responses^2,3^ and cellular signaling^4,5^. The proteins are also widely used as diagnostic tumor markers^4^. Keratin 1 and its heterodimer partner keratin 10 are the major keratins in the suprabasal keratinocytes of epidemis^6,7,8^, which is a hallmarker for keratinocyte differentiation^9^. Increasing evidence has suggested new biological functions of keratins, e.g., keratin 1 is also an integral component of the multiprotein kininogen receptor of endothelial cells^10,11^. Therefore, it is of both theoretical and clinical significance to further elucidate the biochemical and biophysical properties of keratins.

Human autofluorescence (AF) has shown great promise for non-invasive diagnosis of multiple diseases including diabetes^12^ and cancer^13^. Keratins^14^, together with melanin, NADH and FAD, are major epidermal fluorophores^15^. However, three major questions regarding the AF of keratins have remained unanswered: 1) In addition to its basal AF, can the AF of keratins be induced by such factors as UV? 2) What are the mechanisms underlying keratins’ AF? 3) May the AF of keratins have biomedical applications?

Ultraviolet radiation (UV) is a major cause of various skin diseases including skin cancer^16^. Elucidation of the mechanisms underlying UV-induced skin damage is critical for establishing effective strategies for preventing and treating UV-induced skin diseases. Moreover, there has been no biomarker for non-invasive prediction of UV-induced skin damage. Identification of these biomarkers is of great theoretical and clinical importance. Our current study has provided evidence suggesting that UV-induced keratin 1 proteolysis can lead to increased epidermal AF, which may be used as a biomarker for non-invasive prediction of UV-induced skin damage. More importantly, our study has suggested mechanisms underlying the potential of the UV-induced epidermal AF as a biomarker for predicting UV-induced skin damage: UV-induced keratin 1 proteolysis is a critical pathological event in UV-induced skin damage, which may become a promising target for prevention and treatment of UV-induced diseases.

## Materials and Methods

### Materials

All chemicals were purchased from Sigma (St. Louis, MO, USA) except where noted.

### Animal studies

Male C57BL/6Slac mice, ICR mice, and BALB/cASlac-nu nude mice of SPF grade were purchased from SLRC Laboratory (Shanghai, China). All of the animal protocols were approved by the Animal Study Committee of the School of Biomedical Engineering, Shanghai Jiao Tong University.

### Studies on human Subjects

The human subjects of the study included 8 men and 4 women. The Protocol for Human Subject Studies was approved by the Ethics Committee for Human Subject Studies of Shanghai Ninth Hospital, Shanghai Jiao Tong University School of Medicine.

### Exposures of UV radiation

UVA lamp (TL-K 40W ACTINIC BL Reflector, Philips, Hamburg, Germany), UVB lamp (TL 20W/01 RS NARROWBAND, Philips, Hamburg, Germany), and UVC lamp (TUV 25W /G25 T8, Philips, Hamburg, Germany) were used as the UV sources in our experiments. C57BL/6Slac mice, ICR mice, and BALB/cASlac-nu nude mice at the weight between 18-35 g were used for UVC treatment. C57BL/6Slac mice at the weight between 20-30g were used for UVB or UVA treatment. After the mice were briefly anesthetized with 3.5% (w/v) chloral hydrate (1 ml /100 g), the ears of the mice were exposed to UV lamps. The power densities of UVA and UVB were 3.0±0.1 mW/cm^2^ and 2.1±0.1 mW/cm^2^, respectively, measured by a UVA/UVB detector (ST-513, UVAB, SENTRY OPTRONICS CORP., Taiwan, China). The power density of UVC was 0.55±0.05 mW/cm^2^, measured by a UVC detector (ST-512, UVC, SENTRY OPTRONICS CORP., Taiwan, China). The index fingers of human subjects were exposed to a UVC lamp (TUV 25W /G25 T8, Philips, Hamburg, Germany) at the power density of 2.0±0.1 mW/cm^2^, with the radiation dosage of 2.4±0.1 J/cm^2^.

### SR X-ray exposures

All surgery procedures were performed under chloral hydrate anesthesia. C57 mice were irradiated at beam line station BL13W1 of Shanghai Synchrotron Radiation Facility (SSRF) at the dosages of 10 or 20 Gy, using 10.5 or 30.5 KeV SR X-rays. The ear of the anesthetized mice were exposed to SR X-rays, while other parts of the body were protected by lead sheets, as previously described^17^. The radiation doses of SR X-rays were calculated by the air kerma method, as described previously^17^. An ionization chamber was employed to determine the photon flux of SR X-rays by measuring the ionized electron currents. The photon flux was used to calculate the air kerma at the entrance of the tissues, which was converted to the average radiation dosages.

### Imaging of skin AF

The AF of the ears of the mice were imaged by a two-photon fluorescence microscope (A1 plus, Nikon Instech Co., Ltd., Tokyo, Japan), with the excitation wavelength of 488 nm and the emission wavelength of 500 - 530 nm. The AF was quantified by the following approach: Sixteen spots with the size of approximately 100 X 100 μm^2^ on the scanned images were selected randomly. After the average AF intensities of each layer were calculated, the sum of the average AF intensities of all layers of each spot was calculated, which is defined as the AF intensity of each spot. If the value of average AF intensity of certain layer is below 45, the AF signal of the layer is deemed background noise, which is not counted into the sum.

The spectra of the AF of the mice was determined by using a two-photon fluorescence microscope (A1 plus, Nikon Instech Co., Ltd., Tokyo, Japan). After the imaging, the images were analyzed automatically. The index fingers of human subjects were imaged under a confocal fluorescence microscope (TCS SP5ll, Lecia, Wetzlar, Germany). The excitation wavelength was 488 nm and the emission wavelength was 500 - 550 nm. E ight spots with the size of approximately 200 X 200 μm^2^ on the scanned images were selected randomly, which were used for calculations of the AF intensities of the fingers.

Two hrs after administration with E-64, the ears of C57 mouse were irradiated with UVC. Subsequently the ears were collected, which were imaged under a confocal fluorescence microscope (TCS SP5ll, Lecia, Wetzlar, Germany). The excitation wavelength was 488 nm and the emission wavelength was 500-550 nm. The AF was quantified by using the following protocol: Sixteen spots with the size of approximately 100 X 100 μm^2^ on the scanned images were selected randomly. The average AF intensities of the 16 spots were quantified. The average value of the 16 AF intensities was defined as the AF intensity of the sample.

### Histology

Skin biopsies from the ears of the mice were obtained, which were placed immediately in 4% (w/v) paraformaldehyde buffer. After 12-24 hrs, paraffin embedding procedure was conducted on the samples. Hematoxylin / Eosin staining was performed according to the manufacturer’s protocol (Beyotime, Haimen, Jiangsu Province, China).

### Western blot assays

The lysates of the skin were centrifuged at 12,000 *g* for 20 min at 4°C. The protein concentrations of the samples were quantified using BCA Protein Assay Kit (Pierce Biotechonology, Rockford, IL, USA). As described previously^17^, a total of 50 μg of total protein was electrophoresed through a 10% SDS-polyacrylamide gel, which were then electrotransferred to 0.45 μm nitrocellulose membranes (Millipore, CA, USA). The blots were incubated with a monoclonal Anti-Cytokeratin 1 (ab185628, Abcam, Cambridge, UK) (1:4000 dilution) or actin (1:1000, sc-58673, Santa Cruz Biotechnology, Inc., Dallas, TX, USA) with 0.05% BSA overnight at 4°C, then incubated with HRP conjugated Goat Anti-Rabbit IgG (H+L) (1:4000, Jackson ImmunoResearch, PA, USA) or HRP conjugated Goat Anti-mouse IgG (1:2000, HA1006, HuaBio, Zhejiang Province, China). An ECL detection system (Thermo Scientific, Pierce, IL, USA) was used to detect the protein signals. The intensities of the bands were quantified by densitometry using Image J.

### Immunofluorescence

Ten μm paraffin-sections of skin were obtained by a Leica Cryostat, mounted onto poly-L-lysine coated slides and stored at room temperature. The skin sections were incubated in Xylene three times. The sections were then sequentially dehydrated in 100% EtOH, 95% EtOH and 70% EtOH. After two washes with PBS, the sections were blocked in 10% goat serum for 1 hr, which were then incubated in monoclonal Anti-Cytokeratin 1 (ab185628, abcam, Cambridge, UK) (1:1000 dilution), containing 1% goat serum at 4 °C overnight. After three washes in PBS, the sections were incubated with Alexa Fluor 647 goat anti-rabbit lgG (1:1000 dilution) (Invitrogen, CA,USA) for 1 hr in darkness at RT, followed by staining in 0.2% DAPI solution (Beyotime, Haimen, Jiangsu Province, China) for 5 min. Subsequently the sections were mounted in Fluorescence Mounting Medium (Beyotime, Haimen, Jiangsu Province, China). To compare the intensity of fluorescence in each sample, at least three randomly picked fields in each section were photographed under a Leica microscope.

### Laser-based delivery of keratin 1 siRNA into mouse skin

Male C57BL/6SlacMice were briefly anesthetized with 3.5% (w/v) chloral hydrate (1 ml / 100 g). After exposure to laser, the ears of the mouse were transfected with siRNA using Lipofectamine 3000 following the manufacturer’s instructions (Thermo Fisher Scientific, Waltham, MA, USA). The sequences of the mouse keratin siRNA were CUCCCAUUUGGUUUGUAGCTT and UGACUGGUCACUCUUCAGCTT (GenePharma, Shanghai, China).

### Statistical analyses

All data are presented as mean + SEM. Data were assessed by one-way ANOVA, followed by Student–Newman–Keuls *post hoc* test. *P* values less than 0.05 were considered statistically significant.

## Results

### 1) UVC dose-dependently induced increases in the epidermal AF of mouse ears

We determined the effects of UVC exposures on the skin AF of the ears of C57 mice, showing that 0.33, 0.66 and 0.99 J/cm^2^ UVC led to increases in the green AF of the skin (excitation wavelength is 488 nm, and emission wavelength is 500 - 530 nm), assessed at 0, 1, 6, or 24 hrs after the UVC exposures (**Figure 1A**). Quantifications of the AF showed that UVC dose-dependently induced significant increases in the skin AF of the ears (**Figure 1B**). The AF intensities at both 1 and 6 hrs after the UVC exposures had virtually linear relationships with the UVC dosages: r_1h_ = 0.9736 and r_6h_ = 0.9736 for the correlation between the UVC dosages and the average intensities of the green AF at 1 hr and 6 hr after the UVC exposures, respectively. The increased AF remained unchanged for at least 24 hrs after the UVC exposures, except that the AF induced by 0.99 J/cm^2^ UVC was mildly decreased at 24 hrs after the UVC exposures, compared with the AF at 6 hrs after the UVC exposures (**Figure 1B**).

**Figure 1.**
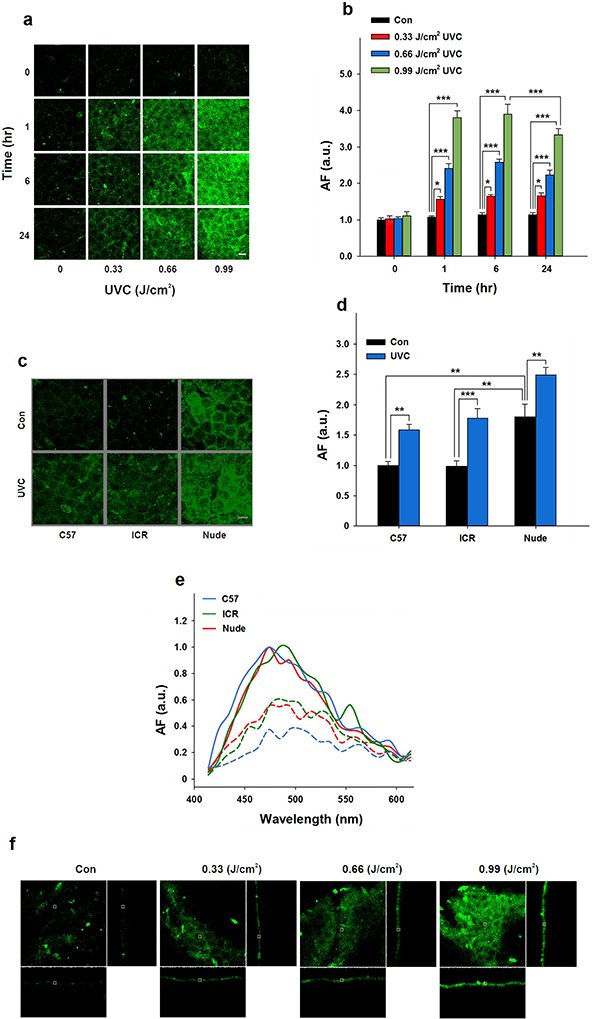
UVC dose-dependently induced increases in the epidermal AF of mouse ears. (A) Exposures of the skin to 0.33, 0.66 or 0.99 J/cm^2^ UVC led to dose-dependent increases in the skin AF of the ears of C57 mice, assessed at 0, 1, 6 or 24 hrs after the UVC exposures. Excitation wavelength = 488 nm and emission wavelength = 500 - 530 nm. Scale bar = 20 μm. (B) Quantifications of the AF showed that each dose of UVC induced significant increases in the skin AF of the ears of C57 mice. *, *P* < 0.05; ***, *P* < 0.001. N = 10 – 13. (C) Exposures of the skin of the ears of ICR and nude mice to UVC also led to increases in the green AF. The AF was assessed at 3 – 6 hrs after the UVC exposures. Scale bar = 20 μm. (D) Quantifications of the AF showed that UVC induced significant increases in the skin AF of the ears of ICR and nude mice. **, *P* < 0.01; ***, *P* < 0.001. N = 4 – 5. (E) The spectra of the UVC-induced AF of C57, ICR and nude mice were similar. The spectra of the epidermal AF of three mice of each strain were determined. The data are the representative of the spectrum of one mice of each strain. (F) The distribution of the AF exhibited the characteristic structure of suprabasal epidermal cells (spinous cells). The AF was originated from the layer of the epidermis which is approximately 10 – 20 μm from the outer layer of the stratum corneasum. The images of XY axis (square), YZ axis (76 μm in length, left column) and XZ axis (76 μm in length, bottom column) were shown.

Exposures of the skin of the ears of nude mice and ICR mice (albino mice) to UVC also led to increases in the green AF (**Figures 1C**). Quantifications of the AF indicated that UVC induced significant increases in the skin AF of the ears, assessed at 3 - 6 hrs after UVC exposures (**Figure 1D**). Using two-photon fluorescence microscopy, we further found that the spectra of the AF of these strains of mice were similar, reaching maximal AF intensity at 470 - 500 nm when the excitation wavelength was 800 nm in two-photon fluorescence microscope (**Figure 1E**).

It is established that UVC can dose-dependently induce skin damage^18^. We determined the UVC-induced skin damaged by H & E staining, showing that the UVC exposures led to significant decreases in the number of epidermal stromal cells (**Supplemental Figures 1A and 1B**) and incrassation of ear (**Supplementary Figures 1C and 1D**) at 72 hrs after the UVC exposures, but not at 24 hrs after the exposures. The UVC-induced AF intensities are positively correlated with the decrease in the number of epidermal stromal cells (**Supplementary Figures 1E**).

In order to investigate the locations of the UVC-induced green AF signals, we studied the orthographic images of the skin of C57 mouse ears. The UVC-induced increases in the AF mainly occurred only at certain layer of the skin, which is approximately 10 - 20 μm from the outer surface of the epidermis (**Figure 1F**). Our H&E staining of the skin of C57 mouse ears indicated that the thickness of the epidermis of the mouse ears is approximately 25 - 30 μm, while the thickness of the stratum corneasum is less that 3 μm (**Supplementary Figures 1A**). Therefore, the AF signals appear to be located between the granular cell layer and the basal cell layer of the epidermis. The spatial distribution of the UVC-induced AF is distinctly polyhydral (**Figure 1a**), exhibiting the characteristic structure of the epidermal cells at the spinous layer of the epidermis^19–21^.

### 2) UVB and UVA dose-dependently induced increases in the epidermal AF of mouse ears

Exposures of the ears of C57 mice to 1.3 and 2.6 J/cm^2^ UVB also led to dose-dependent increases in the green AF, assessed at 24 hrs after the exposures (**Fig. 2A and 2B**). The AF intensities are positively correlated with UVB intensities: r_6h_ = 0.9870 and r_24h_ = 0.9999 for the correlation between the UVB dosages and the average intensities of the green AF at 6 hr and 24 hr after the UVB exposures, respectively. Exposures of the ears of C57 mice to UVA at the dosage of 5.4 J/cm^2^ led to a significant increase in the AF, assessed at 24 hrs after the exposures (**Figure 2C and 2D**). The AF intensities are positively correlated with UVA intensities: r24h = 0.9675 for the correlation between the UVA dosages and the average intensities of the green AF at 24 hr after the UVA exposures. In contrast, irradiation of the ears of C57 mice by SR X-ray did not induce any significant increases in the AF (**Supplemental Figure 2A and 2B**). We also determined the spectra of UVC-, UVB-, and UVA-induced AF by two-photon fluorescence microscopy, showing that the spectra of UVC-, UVB-, and UVA-induced AF were similar, reaching its peak at 470 - 480 nm when the excitation wavelength was 800 nm (**Figure 2E**).

**Figure 2.**
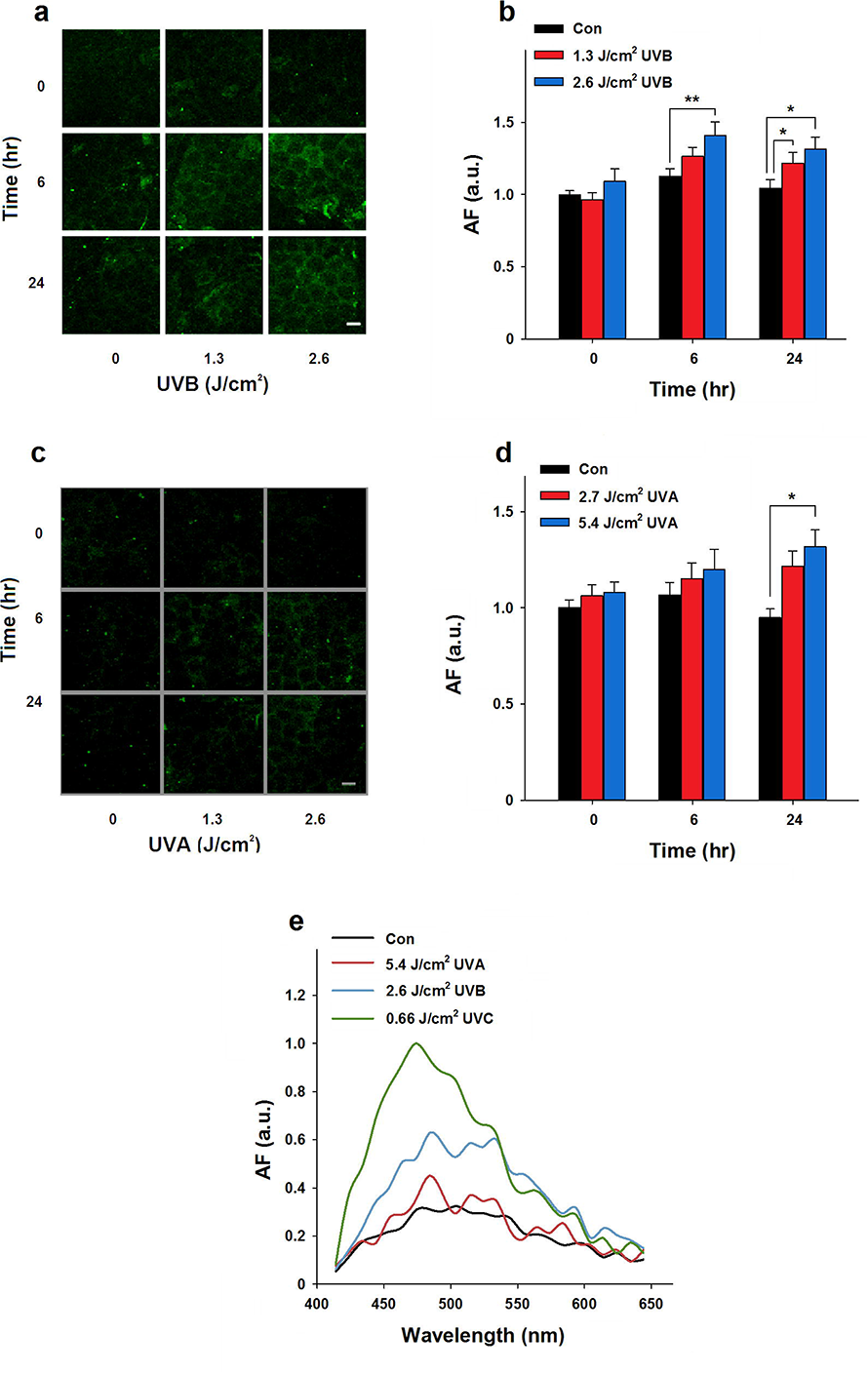
UVB and UVA induced increases in the epidermal AF of mouse ears. (A) Exposures of the ears of C57 mice to 1.3 and 2.6 J/cm^2^ UVB led to increases in the green AF, assessed at 24 hrs after the exposures. Scale bar = 20 μm. (B) Quantifications of the AF showed that each dose of UVC induced significant increases in the skin AF of the ears of C57 mice. *, *P* < 0.05; **, *P* < 0.01. N = 10 – 13. (C) Exposures of the ears of C57 mice to UVA at the dosage of 5.4 J/cm^2^ led to increases in the green AF, assessed at 24 hrs after the exposures. Scale bar = 20 μm. (D) Quantifications of the AF showed that exposures of the ears of C57 mice to UVA at the dosage of 5.4 J/cm^2^ led to a significant increase in the green AF, assessed at 24 hrs after the exposures. *, *P* < 0.05. N = 11-15. (E) The spectra of the UVC-, UVB- and UVA-induced AF of C57 mouse ears were similar. Three C57 mice were used for determining the spectra of the UVC-, UVB- and UVA-induced AF, respectively. Each line in the figure is the representative spectrum of one mouse that was exposed to certain type of UV radiation.

### 3) UVB and UVC induced cysteine protease-mediated keratin 1 proteolysis of C57 mouse ears

The spectrum of the UV-induced AF is highly different from those of FAD, melanin and NADH^22,23^, thus excluding the possibility that NADH, FAD and melanin are the molecules responsible for the UV-induced AF. Our observation that UVC can induce significant increases in the AF from the skin of ICR mice (**Figures 1E**), a strain of albino mice that are melanin deficient^24^, has further excluded the possibility that melanin is the molecule responsible for the AF. There is evidence implicating that keratin 1 may be the molecule responsible for the UV-induced AF: 1) The spectrum of the UV-induced AF (**Figure 1E**) matches well with that of the basal fluorescence of keratins^25^; and 2) the distribution of the UV-induced AF exhibited the characteristic structure of suprabasal epidermal cells (**Figure 1A**), while keratin 1 is a major form of keratin in suprabasal epidermal cells^6,7,8^.

It is noteworthy that under the normal denaturing conditions for Western Blot analysis on the samples prepared from the C57 mouse ears, the antibody of keratin 1 recognizes two bands at 123 kDa and 67 kDa, respectively (**Figure 2A**). We also found that the ratios between the intensity of 67 kDa band and the intensity of 123 kDa band vary from 0 – 1/4 in all of our Western blot analyses (data not shown). It is established that keratin 1 and keratin 10 form heterodimers^26^. Thus the 123 kDa band could be the keratin 1 / keratin 10 heterodimers. We also applied proteomics approach to determine the contents of the band from the Western blot at the position of 123 kDa, showing that both keratin 1 and keratin 10 exist in the band at similar concentrations (data not shown). These observations support the notion that keratin 1 exists in the forms of either keratin 1 monomers or keratin 1 / keratin 10 heterodimers.

Our study showed that 30 min after the ears of C57 mice were exposed to 0.33 J/cm^2^ or 0.66 J/cm^2^ UVC, UVC induced degradation of the keratin 1 at 123 kDa (**Fig. 3A**). Quantifications of the Western blot indicated that UVC induced significant degradation of the keratin 1 at 123 kDa (**Fig. 3B**). UVB also induced degradation of the keratin 1 at 123 kDa at 2 or 6 hrs after the ears of C57 mice were exposed to 2.6 J/cm^2^ UVB (Fig. 3C). Quantifications of the Western blot indicated that UVB time-dependently induced significant degradation of the keratin 1 at 123 kDa (**Fig. 3D**).

**Figure 3.**
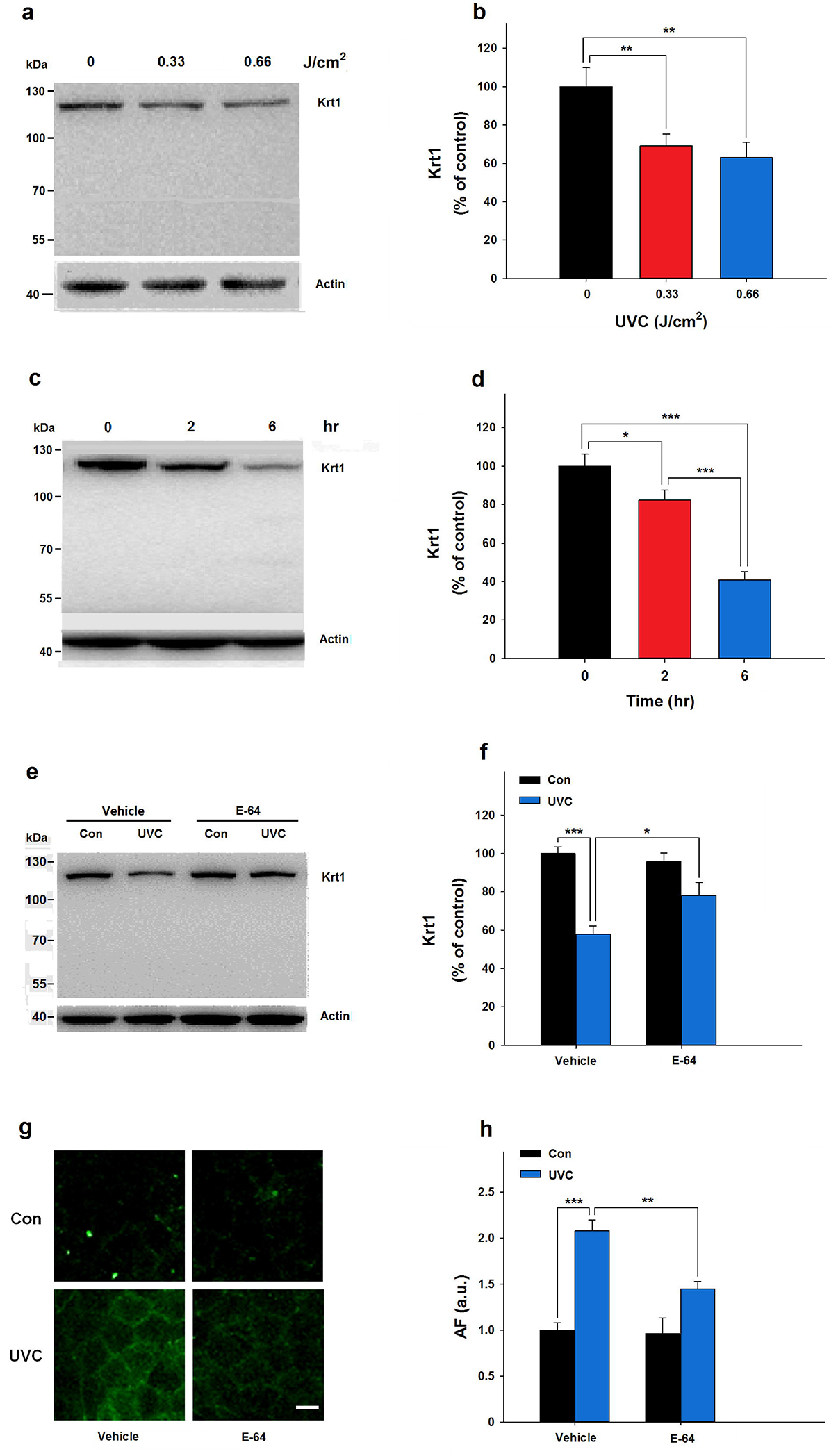
Both UVB and UVC induced cysteine protease-mediated keratin 1 proteolysis of the skin of C57 mouse’s ears. (A) Western blot assays showed that exposures of 0.33 or 0.66 J/cm^2^ UVC dose-dependently induced degradation of keratin 1, assessed at 1 hrs after the UVC exposures. (B) Quantifications of the Western blot showed that UVC induced significant degradation of keratin 1. **, *P* < 0.01. N = 12 – 15. (C) Western blot assays showed that exposures of 2.6 J/cm^2^ UVB induced degradation of keratin 1, assessed at 2 or 6 hrs after the UVB exposures. (D) Quantifications of the Western blot showed that UVB induced significant, time-dependent keratin 1 degradation at both 2 or 6 hrs after the UVB exposures. *, *P* < 0.05, ***, *P* < 0.001. N = 6. (E) Western blot assays showed that administration of E-64, a broad-spectrum cysteine protease inhibitor, led to attenuation of UVC-induced keratin 1 degradation, assessed at 1 hr after the UVC exposures. (F) Quantifications of the Western blot showed that administration with E-64 led to significant attenuation of UVC-induced keratin 1 degradation, assessed at 1 hr after the UVC exposures. *, *P* < 0.05; ***, *P* < 0.001. N = 4 – 6. (G) Administration of E-64 led to attenuation of 0.66 J/cm^2^ UVC-induced green AF of the skin of mouse’s ears, assessed two hrs after the UVC exposures. (H) Quantifications of the AF showed that administration with E-64 led to significant attenuation of UVC-induced increase in the epidermal AF. **, *P* < 0.01; ***, *P* < 0.001. N = 4 – 5.

We applied various protease inhibitors to test our hypothesis that the UVC-induced keratin 1 degradation is mediated by certain proteases. Our *in vivo* studies showed that E-64, a broad-spectrum cysteine protease inhibitor, significantly attenuated UVC-induced decrease in keratin 1 in the ears of C57 mouse (**Figs. 3E and 3F**). The administration of E-64 also led to significant attenuation of the UVC-induced increase in the epidermal AF (**Figs. 3G and 3H**).

### 4) UVC-induced keratin 1 proteolysis mediates the UVC-induced increases in the epidermal AF of mouse ears

We applied laser-based technology to deliver keratin 1 siRNA into the skin to determine the role of keratin 1 in the UVC-induced AF. We found that the laser treatment led to marked increases in the fluorescent signal of Cy5-labelled siRNA inside the skin, compared with that of Cy5-labelled siRNA signal without the laser treatment (data not shown), suggesting the effectiveness of the siRNA delivery. We conducted Western blot assays to determine the effects of keratin 1 siRNA on the keratin 1 levels in the ears. Our Western blot assays on keratin 1 levels showed that the laser-based keratin 1 siRNA administration led to a significant decrease in the keratin 1 levels of the ears (**Figures 4A and 4B**). Immunostaining of the skin also showed that the laser-based keratin 1 siRNA administration led to decreased keratin 1 levels in the epidermis (**Figures 4C**). The keratin 1 siRNA administration was shown to significantly decrease the UVC-induced AF (**Figures 4D and 4E**).

**Figure 4.**
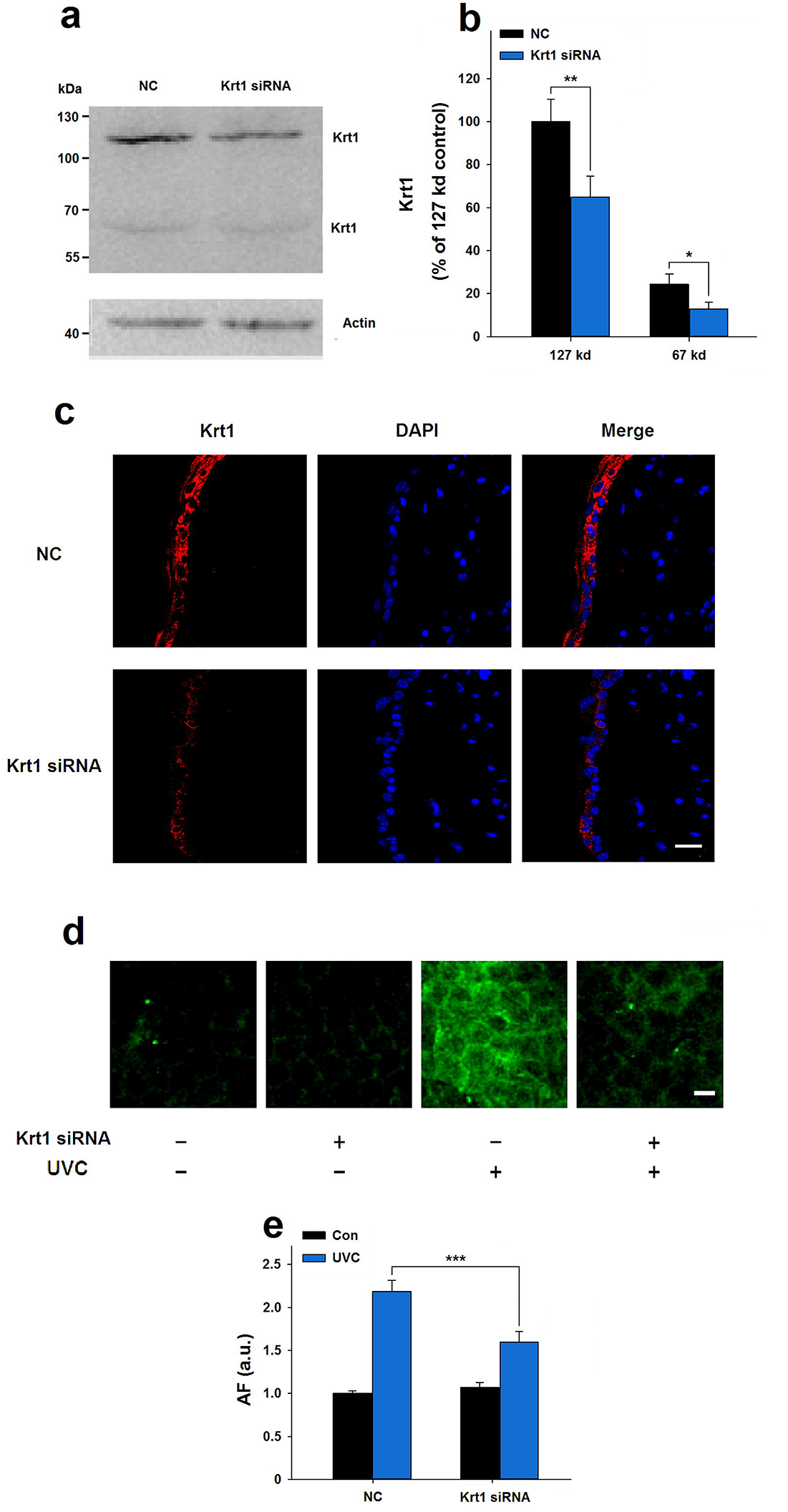
Keratin 1 mediates the UVC-induced increases in the AF of mouse epidermis. (A) Western blot assays showed that the laser-based keratin 1 siRNA administration led to decreases in the keratin 1 levels in the ears of C57 mice. (B) Quantifications of the Western blot showed that the laser-based keratin 1 siRNA administration led to significant decreases in the keratin 1 levels in the ears. *, *P* < 0.05; **, *P* < 0.01. N = 6 – 7. (C) Immunostaining of the skin showed that the laser-based keratin 1 siRNA administration led to an obvious decrease in the keratin 1 levels in the epidermis. (D) The keratin 1 siRNA administration led decreases in the UVC-induced epudermal AF of the ears of C57 mice. (E) Quantifications of the AF indicated that the laser-based keratin 1 siRNA administration led to significant decreases in the UVC-induced AF. ***, *P* < 0.001. N = 6 – 10.

### 5. UVC induced increased skin AF of human subjects

We determined the effect of UVC exposures on the skin AF of human subjects. Exposures of human skin to 2.4 J/cm^2^ UVC led to increased skin AF, assessed immediately after the UVC exposures (**Fig. 5A**). The bar graph shows the AF intensities before or after UVC exposures for each subject (**Fig. 5B**). Quantifications of the AF showed that UVC induced significant increases in human skin AF (**Fig. 5C**).

**Figure 5.**
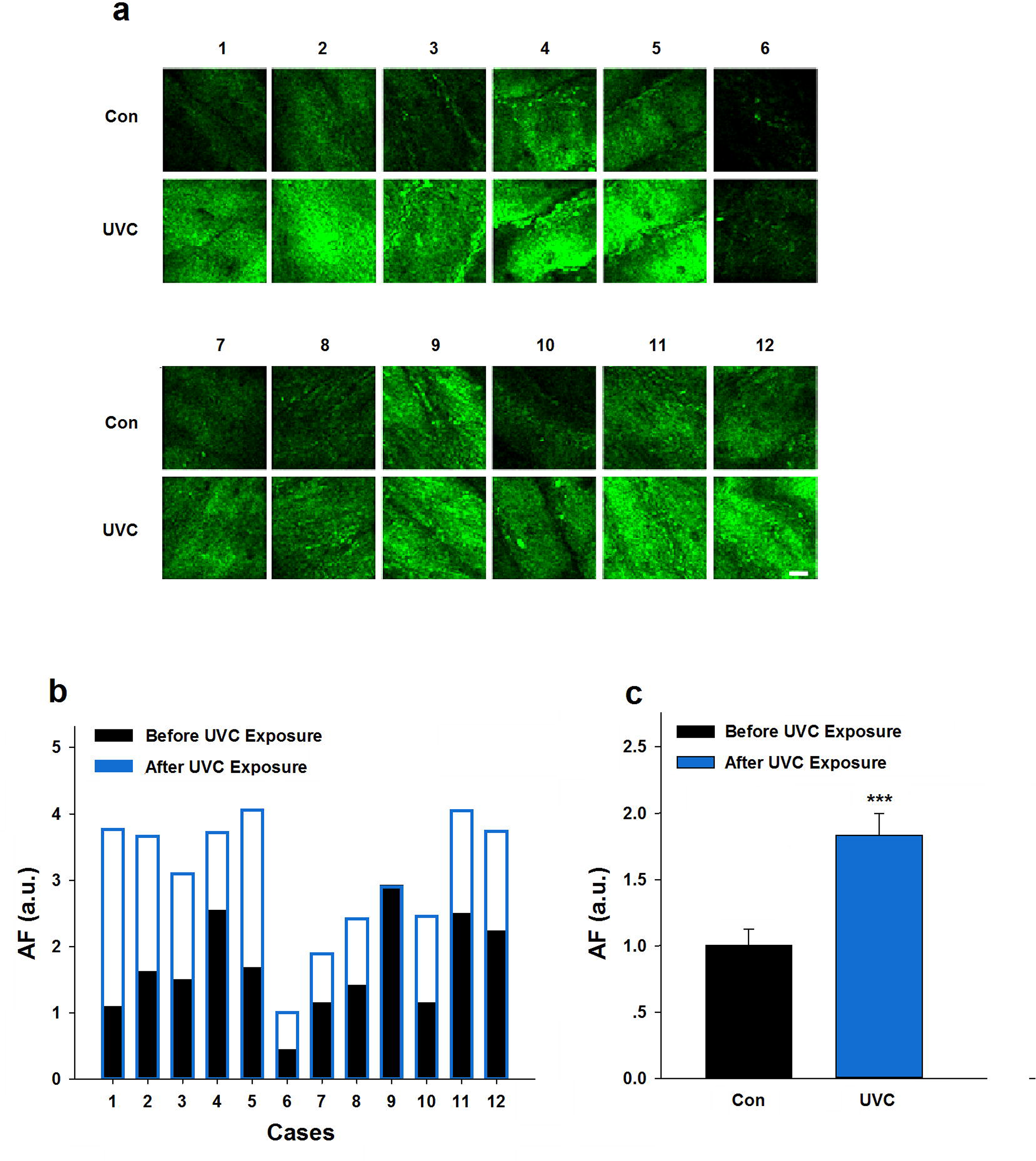
UVC induced increases in the skin AF of human subjects. (A) Exposures of human skin to 2.4 J/cm^2^ UVC led to increased skin AF, assessed immediately after the UVC exposures. Excitation wavelength = 488 nm and emission wavelength = 500 – 530 nm. Scale bar = 100 μm. (B) The bar graph shows the AF intensities before or after the UVC exposures for each subject. (C) Quantifications of the AF showed that UVC induced a significant increase in the AF of human skin. ***, *P* < 0.001. N = 12.

**Figure 6.**
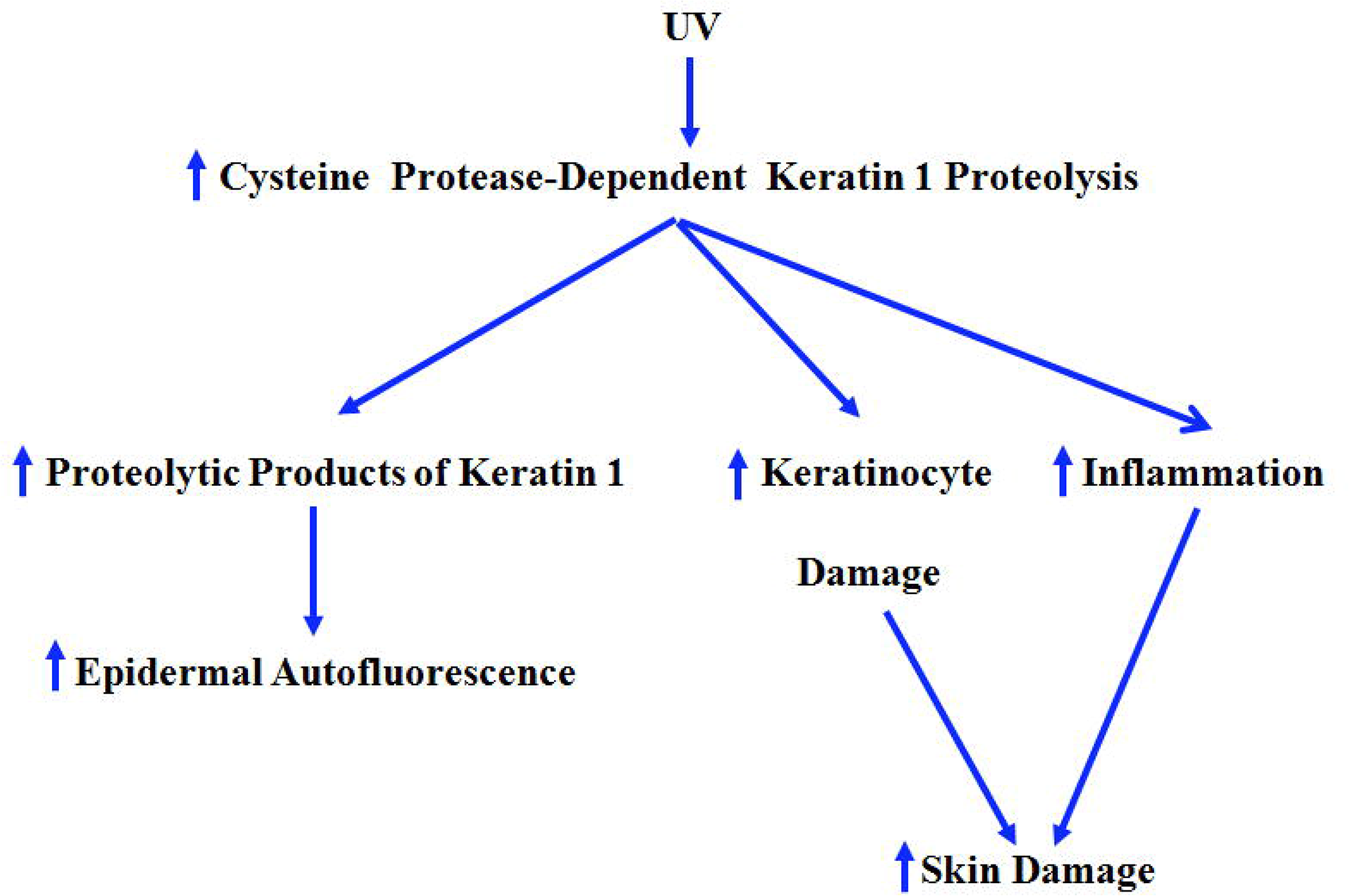
Diagrammatic presentation of the mechanisms underlying the roles of UV-induced keratin 1 proteolysis in UV-induced skin damage.

## Discussion

Our study has provided three lines of evidence indicating that keratin 1 proteolysis is a novel, critical pathological factor in UV-induced skin damage: First, both UVB and UVC can induce rapid and significant keratin 1 degradation; second, prevention of the UV-induced keratin 1 proteolysis by the cysteine protease inhibitor E-64 can significantly attenuate UV-induced skin damage, as indicated by the E-64-produced decrease in the UV-induced epidermal AF; and third, a number of studies have suggested that alterations of keratin 1 can lead to major pathological changes of skin: Keratin 1 plays important roles in intermediate filament formation^1^, inflammatory responses^2,3^, cellular signaling^4,5^ and differentiation^9^; loss of Krt 1 leads to prenatal lethality^2^, in contrast to the mild phenotype observed in Krt 10 null mice^27^; and multiple single mutations of krt 1 lead to several types of cogenital skin disorders^28^. Moreover, it has been reported that approximately 30% amino acid exchanges of keratin 1 are intolerable exchanges.

Our current study has also obtained the following major finding: UV-induced keratin 1 proteolysis can lead to increased epidermal AF, which may be used as a biomarker for predicting UV-induced skin damage. This finding has provided the first evidence indicating that the AF of keratins have significant biomedical applications. Based on this finding, the first device that can predict non-invasively UV-induced skin damage may be developed. This device may be used for not only experimental studies on UV-induced skin damage, but also clinical applications in non-invasive prediction of UV-induced skin damage. In contrast, cell lysis is required in the cyclobutane pyrimidine dimer (CPD)-based approach for predicting UV-induced cellular damage^29^. Moreover, based on our finding, remarkably increased efficacy may be achieved for discovering effective ingredients for decreasing UV-induced skin damage.

Our study has suggested the first mechanism underlying the UV-induced AF of keratin 1: Keratin 1 proteolysis, instead of increased levels of keratin 1, mediates the UV-induced AF. To our knowledge, this finding is the first *in vivo* evidence indicating that proteolysis can lead to increased AF. Because the spectrum of the UV-induced epidermal AF matches well with that of the basal fluorescence of keratins, we speculate that the increased AF might result from either proteolysis-produced increases in the exposures of the fluorescent portions of keratin 1, or proteolysis-produced decreases in the quenching of the basal AF of the keratin 1 fragments.

Our study has also provided first evidence suggesting that the AF of keratins can be induced by UV, in addition to its basal AF. So far most studies regarding applications of AF in disease diagnosis have determined basal AF only, while our current study has suggested that ‘induced AF’ can be used for predicting UV-induced skin damage. By determining both basal AF and ‘induced AF’, significantly richer information may be obtained for medical applications and biomedical research.

In addition to the evidence stated above, our current study has provided the following evidence suggesting that the proteolytic products of keratin 1 may be the origin of the UV-induced AF: First, keratin 1 siRNA-produced decreases in keratin 1 levels led to significant decreases in the UV-induced AF; second, prevention of keratin 1 proteolysis by E-64, a cysteine protease inhibitor, can significantly decrease the UV-induced epidermal AF; and third, our study also showed that overexpression of keratin 1 in B16 cells - a skin cell line - led to a significant increase in the UVC-induced AF (data not shown). Future studies are needed to further demonstrate that the proteolytic products of keratin 1 are the origin of the UV-induced AF.

Our study has applied a laser-based approach with minimal invasiveness to deliver effectively keratin 1 siRNA into the skin, as assessed by Western blotting, immunostaining and AF imaging. To our knowledge, it is the first report that laser-based delivery of siRNA can produce significant biological effects. In contrast, a previous study using laser-based approach showed only that laser can increase the levels of Cy5-labeled siRNA inside the skin, while no evidence was shown that the increase in the siRNA delivery has reached the level that can produce significant biological effects^30^. RNA silencing-based treatment has been a promising strategy for treating skin disorders, while a major technical challenge for this strategy is how to deliver siRNA effectively and non-invasively into the skin^28,31^. Our study has demonstrated that the laser-based approach is an effective and non-invasive approach for *in vivo* delivery of siRNA into the skin.

Several studies have suggested active involvement of keratins in cancer cell invasion and metastasis as well as in treatment responsiveness^4^. Keratin 1 overexpression has also been found in endothelial and mesenchymal tumors^32^. These observations have raised the possibility that novel rapid, non-invasive diagnostic strategies for cancer may be designed on the basis of the UV-induced AF of keratin 1 in the cancer cells. Since keratin 1 share multiple characteristic properties with other Type II keratins^33^, future studies may find similar biological properties of certain Type II keratins as those of keratin 1 found in our study, which may also have significant biomedical applications.

## Acknowledgments

The authors would like to acknowledge the financial support by a Major Research Grant from the Scientific Committee of Shanghai Municipality #16JC1400502 (to W.Y.) and Chinese National Natural Science Foundation Grant #81271305 (to W. Y).

## Legends of Supplemental Figures

**Supplemental Figure 1.** (A) UVC exposures led to obvious decreases in the number of epidermal stromal cells at 72 hrs after the UVC exposures, while no obvious change of the number of epidermal stromal cells was observed at 24 hrs after the UVC exposures. Scale bar = 25 μm. (B) Quantifications of the AF show that UVC exposures led to significant decreases in the number of epidermal stromal cells at 72 hrs after the UVC exposures. *, *P* < 0.05; ***, *P* < 0.001. N = 11 – 16. The UVC exposures did not produce a significant decrease in the number of epidermal stromal cells at 24 hrs after the UVC exposures. N = 5 – 6. (C) UVC exposures led to obvious increases in the incrassation of ear at 72 hrs after the UVC exposures, while no obvious change of the incrassation of ear was observed at 24 hrs after the UVC exposures. Scale bar = 100 μm. (D) Quantifications of the AF show that UVC exposures led to significant increases in the incrassation of ear at 72 hrs after the UVC exposures. *, *P* < 0.05; **, *P* < 0.01. N=12-14. The UVC exposures did not produce a significant change of the incrassation of ear at 24 hrs after the UVC exposures. N = 5 – 6. (E) The UVC-induced AF intensities at 1 hr after the UVC exposures are positively correlated with the UVC-induced decrease in the number of epidermal stromal cells at 72 hrs after the UVC exposures.

**Supplemental Figure 2.** (A) Effects of the exposures of the skin to synchrotron radiation (SR) X-rays at either 10 Gy or 20 Gy on the AF of the ears of C57 mouse. Excitation wavelength = 488 nm and emission wavelength = 500 – 530 nm. Scale bar = 20 μm. (B) Quantifications of the AF did not show that the irradiation of SR X-rays at either 10 Gy or 20 Gy significantly affects the AF of the ears of C57 mouse. N = 4 – 9.

